# Evaluation of host effects on ectomycorrhizal fungal community compositions in a forest landscape in northern Japan

**DOI:** 10.1101/834127

**Authors:** Shunsuke Matsuoka, Yoriko Sugiyama, Ryunosuke Tateno, Shihomi Imamura, Eri Kawaguchi, Takashi Osono

## Abstract

Community compositions of ectomycorrhizal (ECM) fungi are similar within the same host taxa. However, careful interpretation is required to determine whether the combination of ECM fungi and plants is explained by the host preference of ECM fungi or by the influence of neighboring conspecific and/or heterospecific hosts. In the present study, we aimed to evaluate the effects of host species on the ECM community compositions in a forest landscape (~ 10 km) where monodominant forest stands of six ECM host species belonging to three families were patchily distributed. The ECM communities were identified with DNA metabarcoding. A total of 180 ECM operational taxonomic units (OTUs) were detected. The ECM community compositions were primarily structured by host species and families, regardless of the soil environments and spatial arrangements of the sampling plots. In addition, 38 ECM OTUs were detected from particular host tree species. Furthermore, the neighboring plots harbored similar fungal compositions, although the host species were different. The relative effect of the spatial factors on the ECM compositions was weaker than that of host species. Our results suggest that the host preference of ECM fungi is a primary determinant of ECM fungal compositions in the forest landscape.

## Introduction

Ectomycorrhizal (ECM) fungi are symbionts of tree species belonging to the families Fagaceae, Betulaceae, and Pinaceae and represent a dominant group of microorganisms inhabiting temperate and boreal forest floors [1]. ECM fungi play an essential role in plant growth and nutrient cycling by enhancing nutrient and water uptake from soil to their host trees [2]. Since the function or ability of ECM fungi varies from species to species, the community responses of ECM fungi to environmental changes are, therefore, critical for determining and maintaining forest ecosystem processes [3]. So far, various factors, such as host taxa [4], soil properties (e.g., pH) [5], and dispersal limitation [6], have been proposed to affect the compositions of ECM fungal community. For example, environmentally similar or spatially close sites are known to harbor similar ECM fungal communities [5–7]. Practically, the ECM fungal communities in fields are simultaneously affected by these factors. Thus, researchers now try to separate and quantitatively evaluate the effect of each factor on ECM fungal communities and have found the significant effects of host on ECM communities [8–10].

Among these factors, the relationships between host tree species and ECM fungi have been repeatedly tested in variety of regions and/or climatic zones [4, 9–12]. Previous studies have investigated the relatedness of ECM fungal community and host tree species, mainly in single forest stands (mainly < 1 ha) where several host species are mixed, by comparing associated ECM fungi among host individuals in different taxa. These studies have shown that the ECM community compositions are similar within the same host taxa [7, 12]. Such compositional similarities in ECM fungal communities among the same host taxa have often been attributed to the preference of ECM fungi or host for partner species [13, 14].

However, previous studies that investigated the effects of host in a single mixed-forest stand have not necessarily accurately evaluated the host effects owing to some methodological limitations. First, the individuals of the same host species are likely to show clustered distribution in response to the local environmental conditions and past dispersion [15]. In this case, the environmentally similar or spatially close sites tend to harbor similar host communities (i.e., the host community shows correlation with other factors), making it difficult to separate the effects of host and other factors. Second, in mixed-forests, ECM fungal communities are inevitably affected by the surrounding host species. That is, since most fungal spores fall within several meters from sporocarps [16], the spatially closer trees potentially share more inoculums. Furthermore, the same ECM fungal individuals can be shared between adjacent trees via belowground mycelia [17]. Therefore, ECM fungal compositions can be similar among spatially close host trees, regardless of the host taxa [18]. Thus, in most field studies, the effect of each factor has not been fully separated and the effect of host has not been accurately evaluated [8], even though the effects of each factor on ECM fungal communities were evaluated simultaneously.

Among these problems, the correlation between host and other factors and the effects of surrounding host species can be eliminated by conducting surveys in several patchily distributed monodominant forest stands. If the host species has a strong influence, the ECM composition would cluster by host species, regardless of the spatial arrangements of the forest stands. On the other hand, if other environmental factors or spatial distance have stronger effects than the host species, the ECM fungal community compositions should resemble among environmentally similar or spatially closer sites, regardless of the host species.

In the present study, we aimed to evaluate the effect of host trees on ECM fungal community compositions relative to the soil environments and spatial distance at a forest landscape (~ 10 km). Study forests include monodominant forest stands of six ECM host species, including three broad-leaved tree species belonging to the families Fagaceae and Betulaceae and three coniferous species belonging to the family Pinaceae. These forest stands are patchily distributed over a forest. In this setting, we analyzed (1) the effects of the host tree species belonging to three families on the community compositions of ECM fungi, (2) the explanatory power of host tree identities on the ECM compositions relative to other environmental and spatial variables, and (3) the relationships between individual ECM fungal species and host tree species.

## Materials and Methods

### Study sites and Sampling procedure

The study was conducted in the Shibecha branch of Hokkaido Forest Research Station, Field Science Education and Research Center, Kyoto University in the eastern part of Hokkaido Island in northern Japan (1446.8 ha, 43° 22′ N, 144° 37′ E, approximately 25–150 m a.s.l.). The forest area of the station extends approximately 9 km from south to north and approximately 1–3 km from east to west and is surrounded by a pasture. The 30-year mean annual temperature is 6.2 °C, and the 30-year mean annual precipitation of the forest is 1169.7 mm (1981–2010, 43° 19′ N, 144° 36′ E, Kyoto University Forests 2012).

The vegetation of old-growth natural forest is mainly composed of deciduous broad-leaved tree species such as *Quercus crispula* Blume, *Ulmus davidiana* Planchon var. *japonica* (Rehder) Nakai, *Fraxinus mandschurica* Rupr. var. *japonica* Maxim., and *Acer pictum* Thunb. subsp. *dissectum* (Wesm.) H. Ohashi. Pioneer species, such as *Betula platyphylla* Sukaczev and *Alnus hirsuta* (Spach) Turcz. ex Rupr., are patchily distributed on clear-cut areas such as road side and timber yard. The coniferous plantations are monoculture and coniferous species, such as *Larix kaempferi* (Lamb.) Carr., *Abies sachalinensis* F. Schmidt, and *Picea glehnii* F. Schmidt, have been planted from the 1960s to the 1980s in this forest station. *Abies sachalinensis* and *P. glehnii* are common species in the Hokkaido Island, but are not naturally distributed in the forest station. *Larix kaempferi* does not occur naturally on Hokkaido Island, but was introduced from Honshu Island in Japan for afforestation. Approximately 70% of the total area of the forest station is covered by deciduous broad-leaved forests, and the remaining area is occupied by plantation forests in which tree species *L. kaempferi, A. sachalinensis*, and *P. glehnii* cover approximately 14%, 8%, and 2%, respectively.

Six tree species (three broad-leaved and three coniferous species) were targeted as host species. For each host species, three stands (approximately 0.4 ha) where the targeted species dominated as an ECM host species, were chosen as sampling plots (Table 1 and Fig. 1). The latitude, longitude, altitude of each plot and the diameter at breast height (DBH) of individual tree species were recorded. At each plot, we selected 10 host species individuals that had the DBH > 20 cm, and collected a block of surface soil (10 cm × 10 cm × 5 cm from a depth of 5–10 cm), including tree roots within 1 m from each tree trunk. All host tree individuals were spaced at least 3 m apart from each other, thus minimizing the spatial autocorrelation effect of individual ECM fungi [19, 20]. The blocks were kept in plastic bags and frozen at −20°C during the transport to the laboratory. A total of 180 blocks (6 host species × 3 study plots × 10 soil blocks) were used for the study.

**Table 1.**
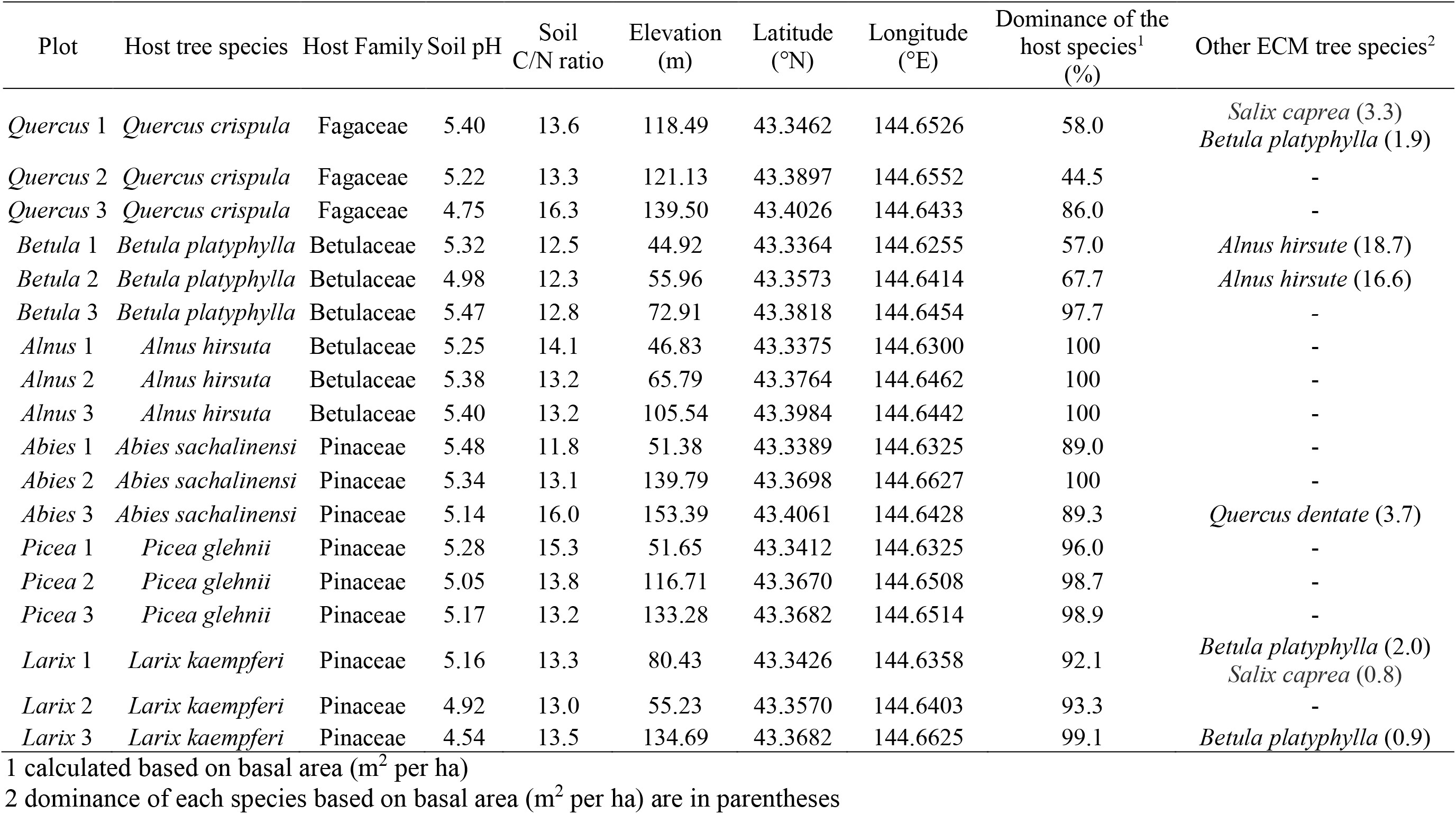
Host tree species and their stand conditions

**Fig. 1.**
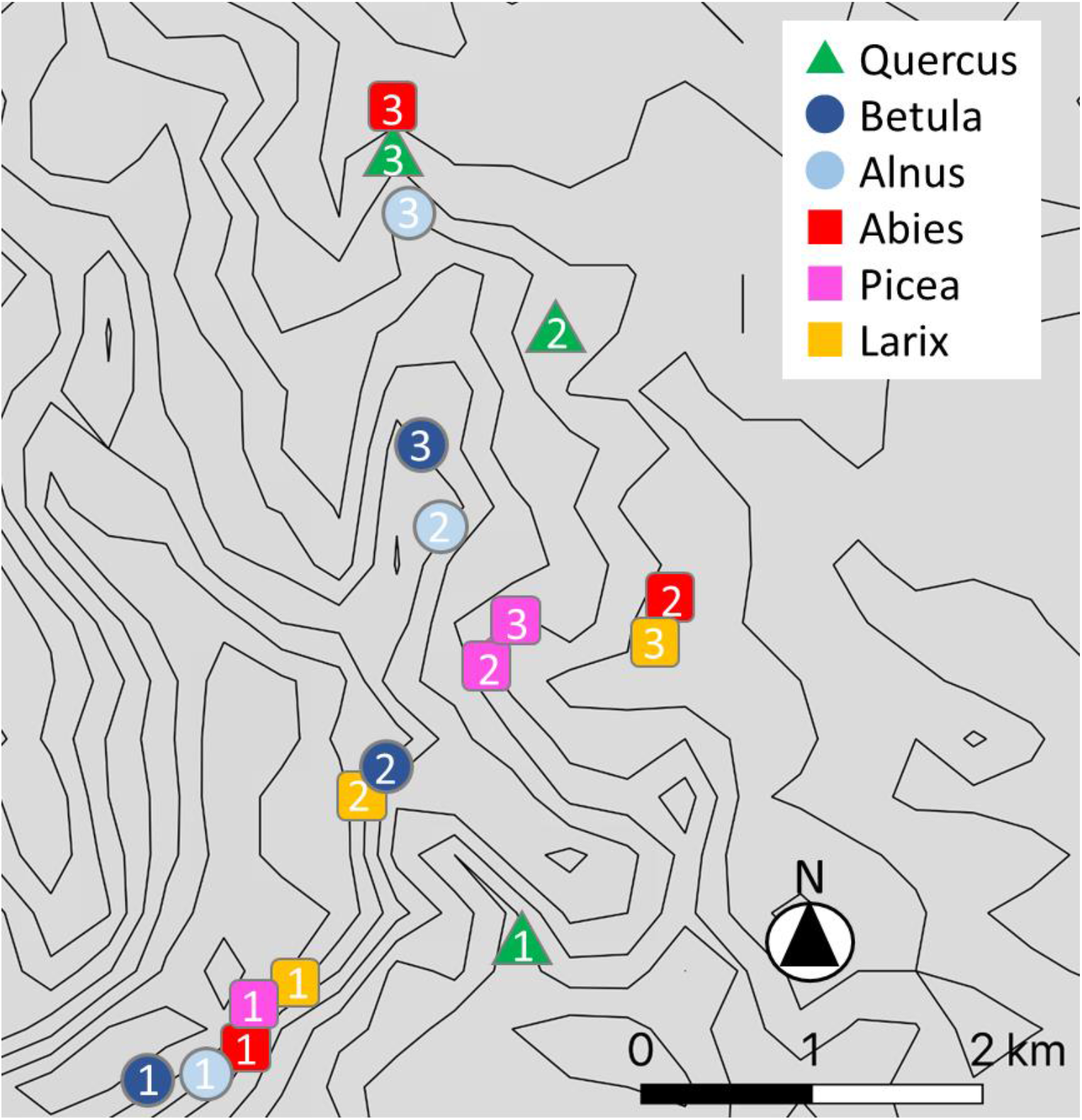
Sampling plots of each tree species. Plot numbers in the symbols are consistent with those listed in Table 1 and Fig. 2.

**Fig. 2.**
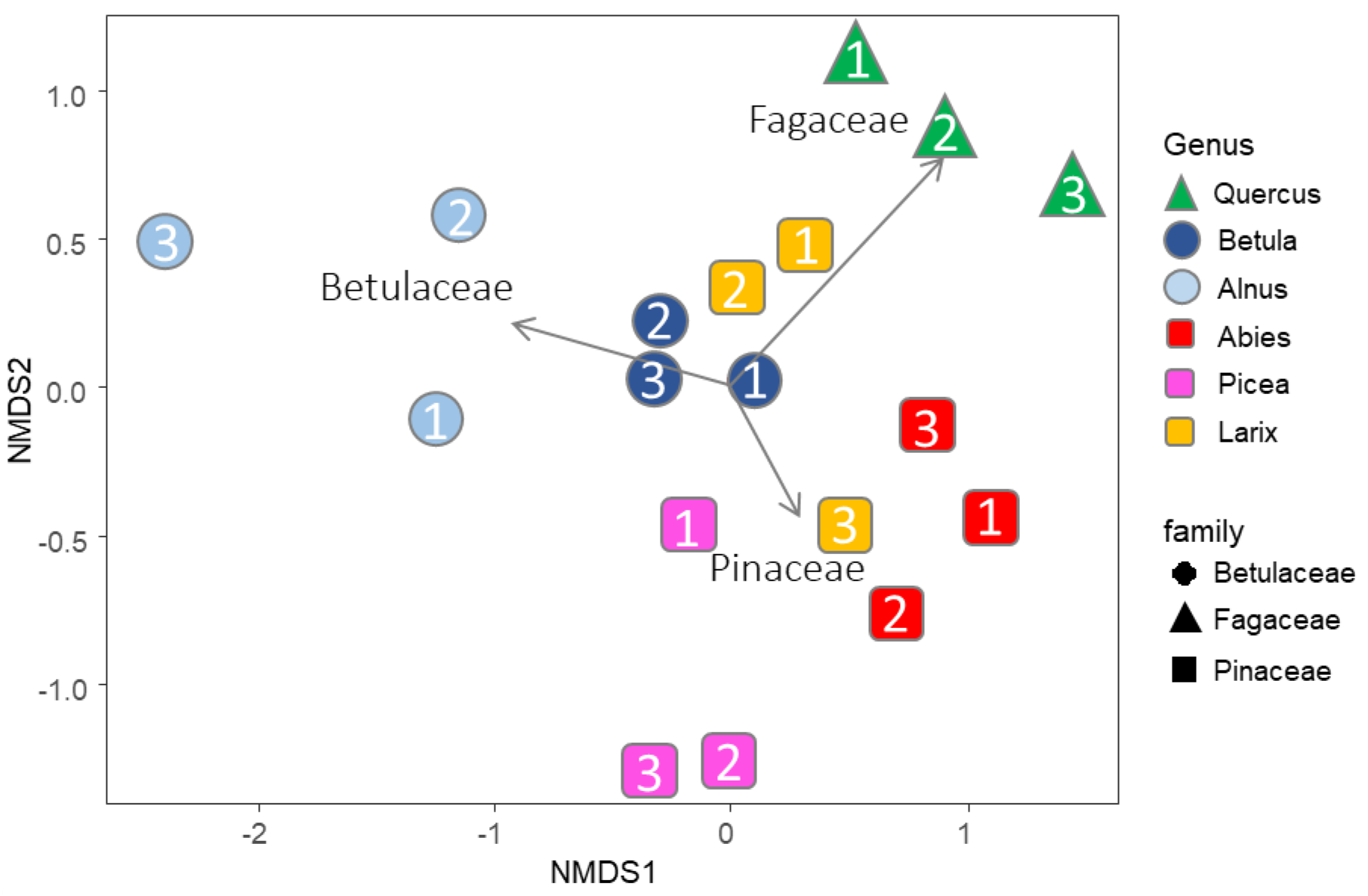
Community dissimilarity among the plots as revealed by nonmetric multidimensional scaling (NMDS) ordination using the Bray-Curtis index (stress value = 0.125). Plot numbers in the symbols are consistent with those listed in Table 1 and Fig. 1.

In the laboratory, fine roots of trees were extracted from the soil samples using a 2-mm mesh sieve and gently washed with tap water to remove the soil particles and debris. In each block, 20 individual root segments (approximately 5 cm in length) were selected, and one root tip (1 to 2 mm in length) was collected from each root segment under a 20X binocular microscope. The 20 root tips resulting from each block were pooled and kept in a tube containing 70% ethanol (w/v) at −20°C. Before extracting DNA, the root tips were washed to remove the small particles on the root surface by 0.005% aerosol OT (di-2-ethylhexyl sodium sulfosuccinate) solution (w/v) and rinsed with sterile distilled water. The root tips were then transferred to the tubes containing cetyltrimethylammonium bromide (CTAB) lysis buffer and stored at −20°C until DNA extraction.

### Soil properties

Mineral soils (0–10 cm in depth) were collected by a soil core sampler (surface area was 20 cm^2^). Five soil core samples were collected at the distance interval of 1.5 m along a straight line from each plot and composited for each plot. The composited soil samples were dried at 70°C for more than 72 hours and sieved through a 4-mm mesh sieve to remove fine roots, pieces of organic matters, and gravels. Total soil N and C were determined by an NC analyzer (Sumigraph NC-900, Sumika Chemical Analysis Service, Ltd., Osaka, Japan), and the soil pH was determined by a pH meter (HORIBA D-51, Horiba, Ltd., Kyoto, Japan) after extraction with deionized water at a dry soil:water ratio of 2:5 (weight/weight).

### DNA extraction, PCR amplification, and pyrosequencing

DNA analysis was generally performed according to methods described by Matsuoka *et al.* [8]. Whole DNA was extracted from root tips in 180 samples using the modified CTAB method described by Gardes and Bruns [21]. For the direct 454 pyrosequencing of the fungal internal transcribed spacer 1 (ITS1) [22], we used a semi-nested PCR protocol. First, the entire ITS region and the 5′-end region of the large subunit were amplified using the fungus-specific primers ITS1F [21] and LR3 [23]. PCR was performed in a 20-μL volume containing 1.6 μL of template DNA, 0.3 μL of KOD FX NEO (TOYOBO, Osaka, Japan), 9.0 μL of 2X buffer, 4.0 μL of dNTPs, 0.5 μL each of the two primers (10 μM), and 4.1 μL of distilled water. The PCR amplification was performed using the following conditions: an initial denaturation step at 94°C for 5 min, followed by 23 cycles of denaturation at 95°C for 30 s, annealing at 58°C for 30 s, and extension at 72°C for 90 s and then a final extension step at 72°C for 10 min. The PCR products were purified using the ExoSAP-IT PCR Product Clean-up Kit (GE Healthcare, Little Chalfont, Buckinghamshire, U.K.). Thereafter, the second PCR targeting the ITS1 region was performed using the ITS1F primer fused with an 8-bp DNA tag [24] and the universal primer ITS2 [25]. The second PCR was performed in a 20-μL volume containing 1.0 μL of template DNA, 0.2 μL of KOD Plus NEO (TOYOBO), 2.0 μL of 10X buffer, 2.0 μL of dNTPs, 0.8 μLeach of the two primers (5 μM), and 13.2 μL of distilled water. The PCR conditions were as follows: an initial denaturation step at 94°C for 5 min, followed by 28 cycles of denaturation at 95°C for 30 s, annealing at 60°C for 30 s, and extension at 72°C for 90 s and a final extension step at 72°C for 10 min. The PCR products were pooled into five libraries and purified using an AMPure Magnetic Bead Kit (Beckman Coulter, California, USA). The pooled products were sequenced in two 1/8 regions using the GS-FLX sequencer (Roche 454 Titanium) at the Graduate School of Science, Kyoto University, Japan. The sequence data were deposited in the Sequence Read Archive of the DNA Data Bank of Japan (accession number: DRA007781).

### Bioinformatics

The bioinformatics analyses were performed using the methods described by Matsuoka *et al.* [8]. Using the 454 pyrosequencing method, 272 358 reads were obtained. These reads were trimmed with sequence quality [26] and sorted into individual samples using the sample-specific tags. The pyrosequencing reads were assembled using Claident pipeline v0.2.2018.05.29 (software available online) [27]. First, the short reads (< 150 bp) and then the potentially chimeric sequences and pyrosequencing errors were removed, using the software programs UCHIME [28] and CD-HIT-OTU [29], respectively. The remaining 204 627 reads were assembled at a threshold similarity of 97%, which is widely used for the fungal ITS region [30], and the resulting consensus sequences represented molecular operational taxonomic units (OTUs). Then singleton OTUs were removed. The consensus sequences of the OTUs are listed in Table S1 (Supporting Information).

To systematically annotate the taxonomy of OTUs, we used Claident v0.2.2018.05.29 [31], which was built upon an automated basic local alignment search tool (BLAST) search using the National Center for Biotechnology Information (NCBI) BLAST+ algorithm [32] and a taxonomy-based sequence identification engine. Using the reference database from the International Nuceotide Sequence Database Collaboration (INSDC) for taxonomic assignment, the sequences homologous to the ITS sequence of each query were fetched, and then the taxonomic assignment was performed based on the lowest common ancestor algorithm [33]. The results of Claident and the number of reads for the OTUs identified are given in Table S1. To screen for the ECM fungi, we referred to the reviews by Tedersoo *et al*. [34] and Tedersoo and Smith [35] and assigned OTUs to the genera and/or families that were predominantly ECM fungi. The resultant ECM fungal OTUs (ECM OTUs) were used for further analyses (see Table S1).

### Data analyses

For all data analyses, the presence or absence of the ECM OTUs was used as the binary data, rather than the quantitative use of read numbers generated from amplicon sequencing [36, 37]. All analyses were performed using the R package, v.3.4.4 [38]. Differences in the sequencing depth of individual samples affect the number of OTUs retrieved, often leading to the underestimation of OTU richness in the samples that had low sequence reads. In our dataset, because the rarefaction curves for all samples reached an asymptote (Fig. S1), we did not conduct rarefaction analysis.

The OTU compositions were compared between plots. First, the presence or absence of ECM OTUs was recorded for each sample. Subsequently, these presence or absence data were merged within the plots, and the incidence data of each OTU for each plot were generated (n = 10 for each plot). The max occurrence of each OTU was 10 for a single plot. To examine the ECM OTU composition, the dissimilarity index of OTU composition between plots was calculated using the Bray-Curtis index in which the incidence of OTUs is considered. In addition, we used the Raup-Crick index in which only the presence or absence of individual OTUs at each plot was used to confirm the robustness of the results, regardless of the other dissimilarity indexes used. The Raup-Crick dissimilarity index is a probabilistic index and is less affected by the species richness gradient among sampling units than the other major dissimilarity indexes, including the Bray-Curtis index [39]. The community dissimilarity of ECM OTUs among plots was ordinated in nonmetric multidimensional scaling (NMDS). The correlation of the NMDS structure with host identity and geographic (i.e., latitude and longitude) and environmental (i.e., elevation, soil pH, and soil C/N ratio) variables was tested by permutation tests (‘envfit’ command in the vegan package, 9999 permutations). Subsequently, in order to investigate whether the dissimilarity of OTU composition is related to the host (species or family) and geographic positions of the plots (latitude and/or longitude), one-way permutational multivariate analysis of variance (PERMANOVA) was conducted.

We used variation partitioning based on the distance-based redundancy analysis (db-RDA, ‘capscale’ command in the vegan package) to quantify the contribution of the host, environmental, and spatial variables to the community structure of ECM fungal OTUs. The relative weight of each fraction (pure, shared, and unexplained fractions) was estimated following the methodology described by Peres-Neto *et al.* [40]. For the distance-based redundancy analysis (db-RDA), we constructed two models including environmental and spatial variables. The detailed methods for variation partitioning are described by Matsuoka *et al.* [8]. First, we constructed environmental models by applying the forward selection procedure (999 permutations with an alpha criterion = 0.05) of Blanchet *et al.* [41]Blanchet et al. (2008). The full models were as follows: [pH + C/N ratio + elevation + host identity]. Thereafter, we constructed the models using spatial variables, which were extracted based on Moran’s Eigenvector Maps (MEM) [42], Borcard et al., 2004. The MEM analysis produced a set of orthogonal variables derived from the geographical coordinates of the sampling locations. The MEM vectors were calculated using the ‘dbmem’ command in the adespatial package. We used the MEM vectors that best accounted for autocorrelation and then conducted forward selection (999 permutations with an alpha criterion = 0.05; the full model contained six MEM variables). Based on these two models, we performed variation partitioning by calculating the adjusted *R*^2^ values for each fraction (Peres-Neto et al., 2006)[40].

To determine which OTU had significantly different frequencies among the host species, an indicator taxa analysis [43] was performed using the “signassoc” function in the “indicspecies” package on the presence or absence data for each sample (n = 180). We used mode = 1 (group-based) and calculated the p-values with 999 permutations after applying Sidak’s correction for multiple testing.

## Results

### Taxonomic assignment

In total, the filtered 204 627 pyrosequencing reads from the 180 samples were grouped into 488 OTUs with 97% sequence similarity (Table S1). Among them, 180 OTUs (53 939 reads) belonged to the ECM fungal taxa, with 169 OTUs belonging to Basidiomycota and 11 OTUs to Ascomycota. Each plot yielded 4 to 35 OTUs (18 OTUs in average). At the family level, 169 OTUs belonged to 20 families, with the common families being Thelephoraceae (75 OTUs, 41.7% of the total number of ECM fungal OTUs) and Russulaceae (26 OTUs, 14.4%). These two families accounted for 41.0 – 69.8 % of the total richness of ECM fungal OTUs in each tree species (Fig. S4).

### Community structures of ECM OTUs

The NMDS ordination showed the separation of ECM OTU composition among plots (Fig. 2, stress value = 0.125). The ordination was significantly correlated with the host species and family (‘envfit’ function; host species, R^2^ = 0.851, P < 0.001; host family, R^2^ = 0.559, P < 0.001), but not with the latitude, longitude, elevation, soil pH, and C/N ratio of the plot (latitude, R^2^ = 0.029, P = 0.795; longitude, R^2^ = 0.052, P = 0.670; elevation, R^2^ = 0.1456, P = 0.308; soil pH, R^2^ = 0.1099, P = 0.4047; soil C/N ratio, R^2^ = 0.0243, P = 0.8334). In the PERMANOVA, both host species and host family significantly affected the ECM composition (host species, F-value = 57.7, R^2^ = 0.960, P < 0.001; host family, F-value = 7.02, R^2^ = 0.484, P < 0.001). In the variation partitioning, only host tree species identity was selected as an environmental variable, and two MEM vectors (MEM 4 and MEM 2) were selected as spatial variables (Fig. 3). The percentages explained by the environmental and spatial fractions were 28.7% and 5.4%, respectively, and no shared fraction was detected between the environmental and spatial variables (Fig. 3). In total, 34.1% of the community variation was explained and the remaining 65.9% was unexplained. Using the Raup-Crick index did not affect the results. The NMDS ordination and results of variation partitioning with the Raup-Crick index are available in Supplementary materials (Figs. S2 and S3).

**Fig. 3.**
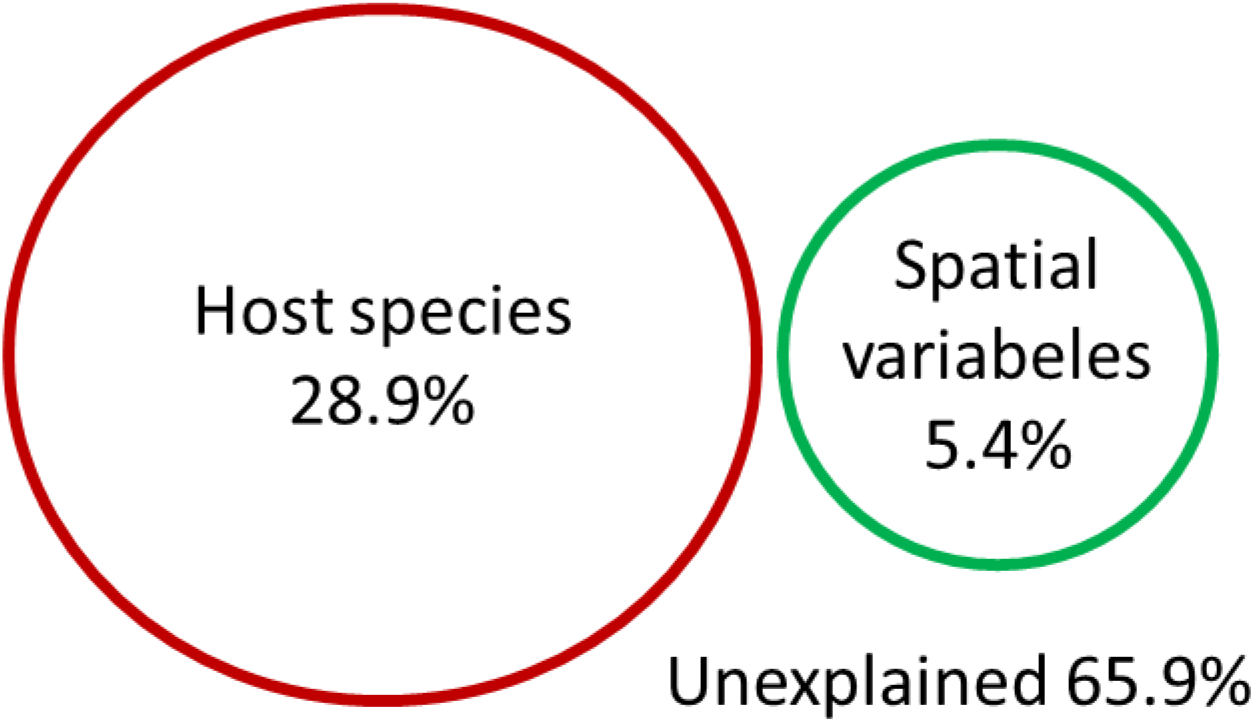
Venn diagram showing the effects of host species and spatial distance on the ectomycorrhizal (ECM) fungal community composition as derived from the variation partitioning analysis. Numbers indicate the proportions of explained variation. No shared fraction between the host species and spatial variables was detected.

The indicator taxa analysis comparing the ECM communities among the host tree species detected significantly different host preferences of 38 OTUs (p < 0.05 after Sidak’s correction, Fig. 4). For each tree, three to nine ECM OTUs showed significantly higher frequencies of occurrence than the other tree species. Different ECM OTUs belonging to the same genus preferred different host tree species. (e.g., OTU_085, OTU_071, and OTU_168 belonging to the genus *Russula* preferred *Quercus, Betula*, and *Picea* tree species as host trees, respectively). In addition, for OTU_109 and OTU_126 belonging to the same ECM fungal species, *Tomentella sublilacina,* the frequently detected host tree species were different, being *Betula* and *Alnus*, respectively.

**Fig. 4.**
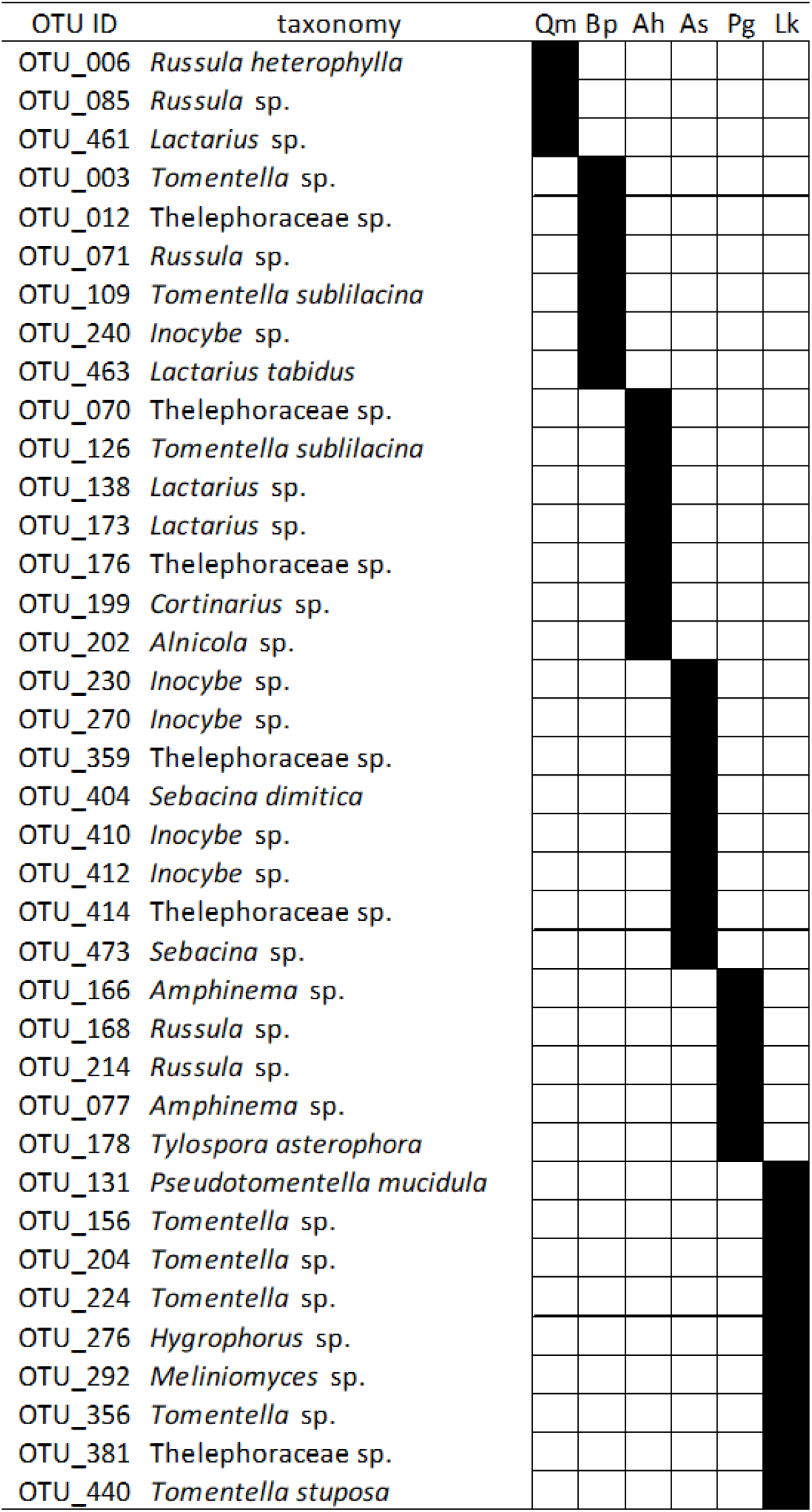
Operational taxonomic units (OTUs) with significantly high detection frequency in particular host tree species. Filled boxes show the combination of ectomycorrhizal (ECM) OTU and tree species with significantly high detection frequency. (P < 0.05 after Sidak’s correction). OTU ID and taxonomy are in accordance with Table S1.

## Discussion

In the present study, we clearly showed the relationships between host species and the ECM fungal community composition by investigating the monodominant forest stands of six ECM host species. We quantitatively evaluated the effect of abiotic environmental and spatial factors on ECM fungal communities in the field, thereby demonstrating the relative importance of host. In our study, the ECM fungal community composition was primarily divided by host species and/or family. In variation partitioning, any part of the fraction explained by host was not shared by the environmental or spatial factors, indicating that we successfully evaluated the pure effect of host in the present study. From this variation partitioning, we could infer the relatively higher importance of host compared to other environmental and spatial factors. In addition, some OTUs showed preference to specific host tree species in our field and could partly contribute to the compositional similarity of ECM fungi within the same host species.

Our results clearly demonstrated that the ECM fungal composition were primarily clustered by host species and phylogeny rather than the soil environments and spatial arrangements of the plots. Similar ECM fungal community composition within the same host species and/or phylogeny has also been detected in other sites and host taxa [4, 8–10, 12]. These similarities in ECM fungal community compositions have been related to the preference of the fungi and/or host tree to partner species, although the exact mechanism of the preference has not been fully revealed. For example, Bogar *et al.* [13] conducted pot experiments with varying symbiotic ability among ECM fungal species, and suggested that plants can discriminate fungal partners and reward more carbohydrates to the fungal species beneficial for the host species. Such selection of the fungal partner by host plants might lead to the different ECM fungal compositions among host species in the field. In addition to these direct interactions between host tree and ECM fungus, environments that the host tree generates (e.g., soil properties) [44, 45] or the interaction with other organisms under particular host species such as soil bacteria or fungus might generate different ECM compositions among host tree species [46].

In our study, the host species has a primary effect on the ECM fungal community composition. However, as the present study is based on the field observation, we cannot infer a causal relationship between a host species and an ECM fungal community. Especially, there is a possibility that unmeasured factors are related to the ECM fungal community composition. For example, in the present study, because the broad-leaved stands are natural forests, the differences in fungi among these stands might partly include the possibility that the fungi and tree species are independently adapted to the same environments [47]. Furthermore, in variation partitioning, 65.9% of the community difference remained unexplained. This unexplained fraction might include the effects of the vegetation of the surrounding area, unexplained environmental factors (e.g., soil organic phosphorus) [48], and drift (i.e., random arrival and extinction) [49].

In our site, the detection of some OTUs was biased to specific host species (Fig. 4). These OTUs might have a high host preference (Fig. 4). Different OTUs belonging to same genus preferred different host tree species. (e.g., OTU_085, OUT_071, and OUT_168 belonging to the genus *Russula* preferred *Quercus, Betula*, and *Picea* tree species as host trees. Moreover, although OTU_109 and OTU_126 were identified as the same species, *Tomentella sublilacina,* the host species frequently associated with these two OTUs were different (*Betula* and *Alnus*, respectively). *Tomentella sublilacina* has been detected from various regions and hosts tree species in the Northern Hemisphere [11, 47], and its preferred host tree species might be different among genotypes and/or habitats. Our results indicate that the degree of preference and the preferred host is different at the fungal species or genotype level, rather than at the genus or family level in our study forests.

Besides host species, the effect of spatial distance on the ECM fungal community composition was detected. This indicates that the ECM fungal compositions become similar at spatially close sites, regardless of the host trees. For example, in the present study, the ECM fungal communities were similar between the *Betula* and *Larix* forests and between the *Abies* and *Larix* forests (Fig. 2). These high similarities of ECM compositions can be partly due to the geographical closeness of the *Betula* 2 and *Larix* 2 plots and between the *Abies* 2 and *Larix* 3 plots (c.a. 100 m, Table 1 and Fig. 1). As factors that lead to such spatial structures at a small spatial scale (c.a. < 100 m), dispersal and colonization limitations can be suggested. Though the dispersal distances of fungal spores are not fully understood, a previous study revealed that most spores fall within several meters from sporocarps [16]. Thus, spatially closer plots potentially share more inoculums. Moreover, in spatially closer plots, the same ECM fungal individuals can be shared between different host species via belowground mycelia. Such sharing of inoculum and/or mycelia might result in the sharing of ECM species between different adjacent tree species [17, 18]. In our study site, for example, OTU_109 was detected both from *Betula* 2 and *Larix* 2. This OTU prefers *Betula* (Fig. 4); therefore, the detection of this OTU from the *Larix* plot might be due to the infections induced by mycelia. As few studies have investigated the distance limitation in such infections via mycelia, elucidating the importance and frequency of these infections to an unpreferred host needs further investigation. Nevertheless, in our results, the ECM fungal communities were shown to be similar between neighboring plots existing within a distance of 100 m, although the host tree species were different. Such spatial structure can hinder the investigation of fungal-plant combinations caused by partner preferences.

In summary, in the present study, we clearly demonstrated that the ECM fungal communities were primarily structured by the species and families of hosts in a forest landscape. Our results further suggest that the preference of fungi and/or host to partner species can primarily structure ECM fungal compositions in the field. In addition, in our study site, the neighboring plots harbored similar fungal communities, though the host species were different, and the effect of the spatial distance on the similarity was also suggested. Therefore, in order to clarify the preferred host species of individual ECM fungi in fields, further studies considering the spatial configuration of the host tree individual and spatial factors are necessary. The mechanisms by which host preference occurs and further observations of relationships between ECM fungal composition and host identities in other host species and environments would be the future research topics.

## Data Accessibility

All data of the 454 sequencing was shared in DRA (Accession number: DRA007781)

## Acknowledgments

We thank the staffs of Hokkaido Forest Research Station, Field Science Education and Research Center, Kyoto University and Celina Sakaguchi for assistance in fieldwork; Miyuki Hirata for assistance in laboratory work; Koichi Ito for useful discussions; and Hirotoshi Sato and Hideyuki Doi for critical comments on the manuscripts.

## Author contributions

SM, TO, and RT designed the study and SM, SI, TO, and RT contributed to field survey and sampling. SM, YS, and EK contributed to molecular experiments. SM and YS analyzed the data and interpreted the results. SM, YS, RT, and TO wrote the initial draft of the manuscript. All other authors critically reviewed the manuscript.

## References

1. Brundrett MC. 2009. Mycorrhizal associations and other means of nutrition of vascular plants: understanding the global diversity of host plants by resolving conflicting information and developing reliable means of diagnosis. Plant Soil. 320, 37–77. doi:10.1007/s11104-008-9877-9

2. Smith S, Read D. 2008. Mycorrhizal Symbiosis. New York: Academic Press.

3. Lilleskov EA, Parrent JL. 2007. Can we develop general predictive models of mycorrhizal fungal community-environment relationships? New Phytol. 174, 250–256. doi:10.1111/j.1469-8137.2007.02023.x

4. Ishida TA, Nara K, Hogetsu T. 2007. Host effects on ectomycorrhizal fungal communities: insight from eight host species in mixed conifer-broadleaf forests. New Phytol. 174, 430–440. doi:10.1111/j.1469-8137.2007.02016.x

5. Toljander JF, Eberhardt U, Toljander YK, Paul LR, Taylor AFS. 2006. Species composition of an ectomycorrhizal fungal community along a local nutrient gradient in a boreal forest. New Phytol. 170, 873–884. doi:10.1111/j.1469-8137.2006.01718.x

6. Peay KG, Schubert MG, Nguyen NH, Bruns TD. 2012. Measuring ectomycorrhizal fungal dispersal: macroecological patterns driven by microscopic propagules. Mol Ecol. 21, 4122–4136. doi:10.1111/j.1365-294X.2012.05666.x

7. Peay KG, Bruns TD. 2014. Spore dispersal of basidiomycete fungi at the landscape scale is driven by stochastic and deterministic processes and generates variability in plant-fungal interactions. New Phytol. 204: 180–191. doi:10.1111/nph.12906

8. Matsuoka S, Mori AS, Kawaguchi E, Hobara S, Osono T. 2016. Disentangling the relative importance of host tree community, abiotic environment and spatial factors on ectomycorrhizal fungal assemblages along an elevation gradient. FEMS Microbiol Ecol. 92, fiw044. doi:10.1093/femsec/fiw044

9. Boeraeve M, Honnay O, Jacquemyn H. 2018. Effects of host species, environmental filtering and forest age on community assembly of ectomycorrhizal fungi in fragmented forests. Fungal Ecol. 36, 89–98. doi:10.1016/j.funeco.2018.08.003

10. Wang Z, Jiang Y, Deane DC, He F, Shu W, Liu Y. 2019. Effects of host phylogeny, habitat and spatial proximity on host specificity and diversity of pathogenic and mycorrhizal fungi in a subtropical forest. New Phytol. 223, 462–474. doi: 10.1111/nph.15786

11. Horton TR, Bruns TD. 1998. Multiple-host fungi are the most frequent and abundant ectomycorrhizal types in a mixed stand of Douglas fir (*Pseudotsuga menziesii*) and bishop pine (*Pinus muricata*). New Phytol. 139, 331–339. doi:10.1046/j.1469-8137.1998.00185.x

12. Tedersoo L, Mett M, Ishida TA, Bahram M. 2013. Phylogenetic relationships among host plants explain differences in fungal species richness and community composition in ectomycorrhizal symbiosis. New Phytol. 199, 822–831. doi:10.1111/nph.12328

13. Bogar L, Peay K, Kornfeld A, Huggins J, Hortal S, Anderson I, et al. 2019. Plant-mediated partner discrimination in ectomycorrhizal mutualisms. Mycorrhiza. 29, 97–111. doi:10.1007/s00572-018-00879-7

14. Molina RM, Horton TR. 2015. Mycorrhizal specificity: Its role in the development and function of common mycelial networks. In: Horton TR., editor. Mycorrhizal Networks. Dordrecht: Springer. pp. 1–39.

15. Punchi-Manage R, Wiegand T, Wiegand K, Getzin S, Gunatilleke CVS, Gunatilleke IAUN. 2014. Effect of spatial processes and topography on structuring species assemblages in a Sri Lankan dipterocarp forest. Ecology. 95, 376–386. doi:10.1890/12-2102.1

16. Galante TE, Horton TR, Swaney DP. 2011. 95% of basidiospores fall within 1 m of the cap: a field-and modeling-based study. Mycologia. 103, 1175–1183. doi:10.3852/10-388

17. Bogar LM, Kennedy PG. 2013. New wrinkles in an old paradigm: neighborhood effects can modify the structure and specificity of *Alnus*-associated ectomycorrhizal fungal communities. FEMS Microbiol. Ecol. 83, 767–777. doi:10.1111/1574-6941.12032

18. Hubert NA, Gehring CA. 2008. Neighboring trees affect ectomycorrhizal fungal community composition in a woodland-forest ecotone. Mycorrhiza. 18, 363–374. doi:10.1007/s00572-008-0185-2

19. Lilleskov EA, Bruns TD, Horton TR, Taylor D, Grogan P. 2004. Detection of forest stand-level spatial structure in ectomycorrhizal fungal communities. FEMS Microbiol. Ecol. 49, 319–332. doi:10.1016/j.femsec.2004.04.004

20. Bahram M, Peay KG, Tedersoo L. 2015. Local-scale biogeography and spatiotemporal variability in communities of mycorrhizal fungi. New Phytol. 205, 1454–1463. doi:10.1111/nph.13206

21. Gardes M, Bruns TD. 1993. ITS primers with enhanced specificity for basidiomycetes - application to the identification of mycorrhizae and rusts. Mol Ecol. 2, 113–118. doi:10.1111/j.1365-294X.1993.tb00005.x

22. Schoch CL, Seifert KA, Huhndorf S, Robert V, Spouge JL, Levesque CA, et al. 2012. Nuclear ribosomal internal transcribed spacer (ITS) region as a universal DNA barcode marker for Fungi. Proc Natl Acad Sci U S A. 109, 6241–6246. doi:10.1073/pnas.1117018109

23. Vilgalys R, Hester M. 1990. Rapid genetic identification and mapping of enzymatically amplified ribosomal DNA from several *Cryptococcus* species. J Bacteriol. 172, 4238–4246. doi:10.1128/jb.172.8.4238-4246.1990

24. Hamady M, Walker JJ, Harris JK, Gold NJ, Knight R. 2008. Error-correcting barcoded primers for pyrosequencing hundreds of samples in multiplex. Nat Methods. 5, 235–237. doi:10.1038/nmeth.1184

25. White TJ, Bruns TD, Lee SB, Taylor JW. 1990. Amplification and direct sequencing of fungal ribosomal RNA genes for phylogenetics. In: Innis MA, Gelfand DH, Sninsky JJ, T.J White, editors. PCR Protocols – a Guide to Methods and Applications. San Diego: CA Academic Press. pp. 315–322.

26. Kunin V, Engelbrektson A, Ochman H, Hugenholtz P. 2010. Wrinkles in the rare biosphere: pyrosequencing errors can lead to artificial inflation of diversity estimates. Environ Microbiol. 12, 118–123. doi:10.1111/j.1462-2920.2009.02051.x

27. Tanabe AS. 2018. Claident. A software distributed by the author. [cited 2019 October]. Available from: https://www.claident.org/

28. Edgar RC, Haas BJ, Clemente JC, Quince C, Knight R. 2011. UCHIME improves sensitivity and speed of chimera detection. Bioinformatics. 27, 2194–2200. doi:10.1093/bioinformatics/btr381

29. Li W, Fu L, Niu B, Wu S, Wooley J. 2012. Ultrafast clustering algorithms for metagenomic sequence analysis. Brief Bioinform. 13, 656–668. doi:10.1093/bib/bbs035

30. Osono T. 2014. Metagenomic approach yields insights into fungal diversity and functioning. In: Sota T, Kagata H, Ando Y et al. (eds). Species Diversity and Community Structure. 1–23. Berlin, Germany: Springer.

31. Tanabe AS, Toju H. 2013. Two new computational methods for universal DNA barcoding: A benchmark using barcode sequences of bacteria, archaea, animals, fungi, and land plants. PLoS ONE. 8, e76910. doi:10.1371/journal.pone.0076910

32. Camacho C, Coulouris G, Avagyan V, Ma N, Papadopoulos J, Bealer K, et al. 2009. BLAST+: architecture and applications. BMC Bioinform. 10, 421. doi:10.1186/1471-2105-10-421

33. Huson DH, Auch AF, Qi J, Schuster SC. 2007. MEGAN analysis of metagenomic data. Genome Res. 17, 377–386. doi:10.1101/gr.5969107

34. Tedersoo L, May TW, Smith ME. 2010. Ectomycorrhizal lifestyle in fungi: global diversity, distribution, and evolution of phylogenetic lineages. Mycorrhiza. 20, 217–263. doi:10.1007/s00572-009-0274-x

35. Tedersoo L, Smith ME. 2013. Lineages of ectomycorrhizal fungi revisited: Foraging strategies and novel lineages revealed by sequences from belowground. Fungal Biol Rev. 27, 83–99. doi:10.1016/j.fbr.2013.09.001

36. Amend AS, Seifert KA, Bruns TD. 2010. Quantifying microbial communities with 454 pyrosequencing: does read abundance count? Mol Ecol. 19, 5555–5565. doi:10.1111/j.1365-294X.2010.04898.x

37. Elbrecht V, Leese F. 2015. Can DNA-based ecosystem assessments quantify species abundance? Testing primer bias and biomass—sequence relationships with an innovative metabarcoding protocol. PLoS ONE. 10, e0130324. doi:10.1371/journal.pone.0130324

38. R Core Team. 2018. R: A language and environment for statistical computing. R Foundation for Statistical Computing, Vienna, Austria. http://www.R-project.org

39. Chase JM. 2010. Stochastic community assembly causes higher biodiversity in more productive environments. Science. 328, 1388–1391. doi:10.1126/science.1187820

40. Peres-Neto PR, Legendre P, Dray S, Borcard D. 2006. Variation partitioning of species data matrices: estimation and comparison of fractions. Ecology. 87, 2614–2625. doi: 10.1890/0012-9658(2006)87[2614:VPOSDM]2.0.CO;2

41. Blanchet FG, Legendre P, Borcard D. 2008. Forward Selection of explanatory variables. Ecology. 89, 2623–2632. doi: 10.1890/07-0986.1

42. Borcard D, Legendre P, Avois-Jacquet C, Tuomisto H. 2004. Dissecting the spatial structure of ecological data at multiple scales. Ecology. 85, 1826–1832. doi:10.1890/03-3111

43. De Cáceres M, Legendre P. 2009. Associations between species and groups of sites: indices and statistical inference. Ecology. 90, 3566–3574. doi:10.1890/08-1823.1

44. Huggins JA, Talbot J, Gardes M, Kennedy PG. 2014. Unlocking environmental keys to host specificity: differential tolerance of acidity and nitrate by *Alnus*-associated ectomycorrhizal fungi. Fungal Ecol. 12, 52–61. doi:10.1016/j.funeco.2014.04.003

45. Ushio M, Aiba S, Takeuchi Y, Iida Y, Matsuoka S, Repin R, et al. 2017. Plant-soil feedbacks and the dominance of conifers in a tropical montane forest in Borneo. Ecol Monogr. 87, 105–129. doi:10.1002/ecm.1236

46. Nakayama M, Imamura S, Taniguchi T, Tateno R. 2019. Does conversion from natural forest to plantation affect fungal and bacterial biodiversity, community structure, and co-occurrence networks in the organic horizon and mineral soil? Forest Ecol Manag. 446, 238–250. doi:10.1016/j.foreco.2019.05.042

47. Miyamoto Y, Sakai A, Hattori M, Nara K. 2015. Strong effect of climate on ectomycorrhizal fungal composition: evidence from range overlap between two mountains. ISME J. 9, 1870–1879. doi:10.1038/ismej.2015.8

48. Kluber LA, Carrino-Kyker SR, Coyle KP, DeForest JL, Hewins CR, Shaw AN, et al. 2012. Mycorrhizal response to experimental pH and P manipulation in acidic hardwood forests. PLoS ONE. 7, e48946. doi:10.1371/journal.pone.0048946

49. Hubbell SP. 2001. The unified neutral theory of biodiversity and biogeography. New York: Princeton University Press.

