## Supporting Information for "Evaluation of host effects on ectomycorrhizal fungal community compositions in a forest landscape in northern Japan"

The following Supporting Information is available for this article:

Table S1 List of all OTUs, the number of their squence reads and conensus sequence, and taxonomic assignments by Claident

Fig. S1 Rarefaction curve for each soil sample (n = 180).

Fig. S2 Community dissimilarity among the plots as revealed by nonmetric multidimensional scaling (NMDS) ordination using the Raup-Crick index (stress value = 0.132). Plot numbers in the symbols are consistent with those listed in Table 1 and Fig. 1.

Fig. S3 Venn diagram showing the effects of host species and spatial distance on the ectomycorrhizal (ECM) fungal community composition as derived from the variation partitioning analysis using the Raup-Crick index. Numbers indicate the proportions of explained variation. No shared fraction between the host species and spatial variables was detected.

Fig. S4 Family level proportions of ECM fungal OTU for each host tree species.

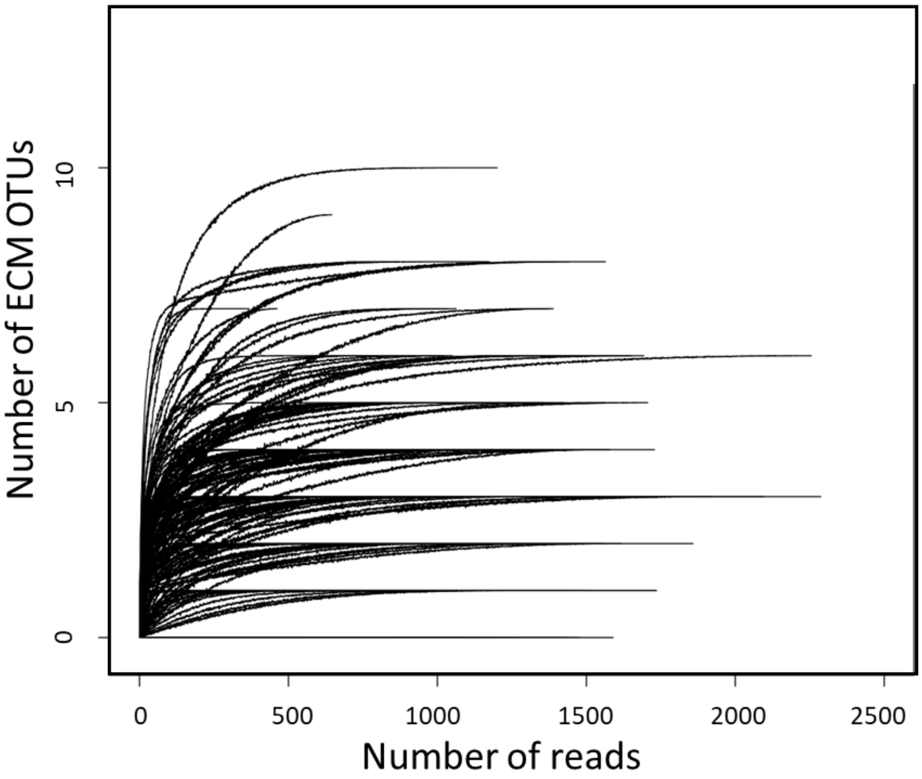


Fig. S1 Rarefaction curve for each soil sample (n = 180).

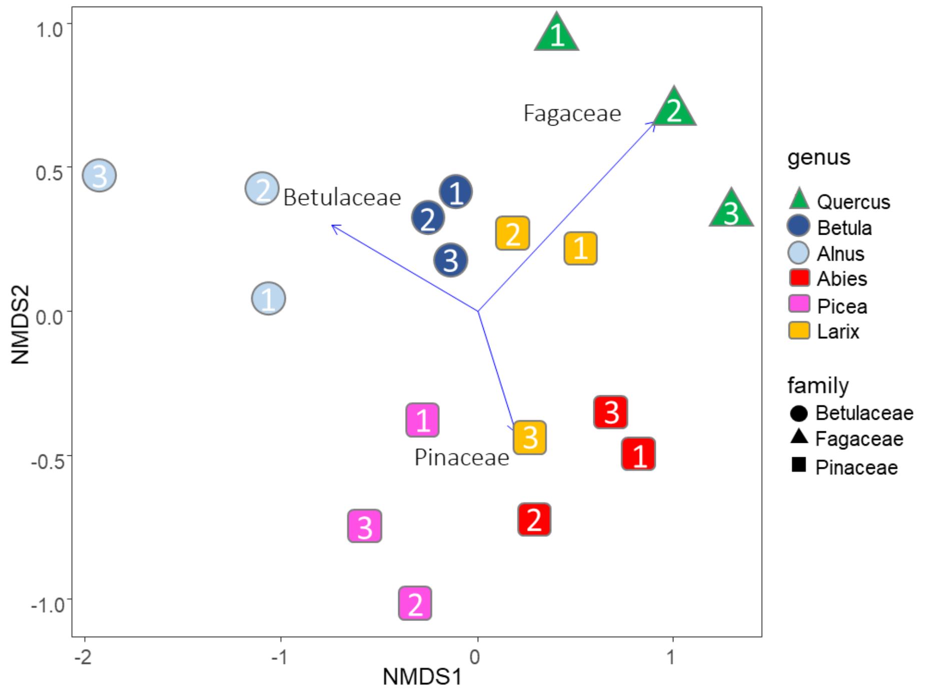


Fig. 2 Community dissimilarity among the plots as revealed by nonmetric multidimensional scaling (NMDS) ordination using the Raup-Crick index (stress value = 0.132). Plot numbers in the symbols are consistent with those listed in Table 1 and Fig. 1.

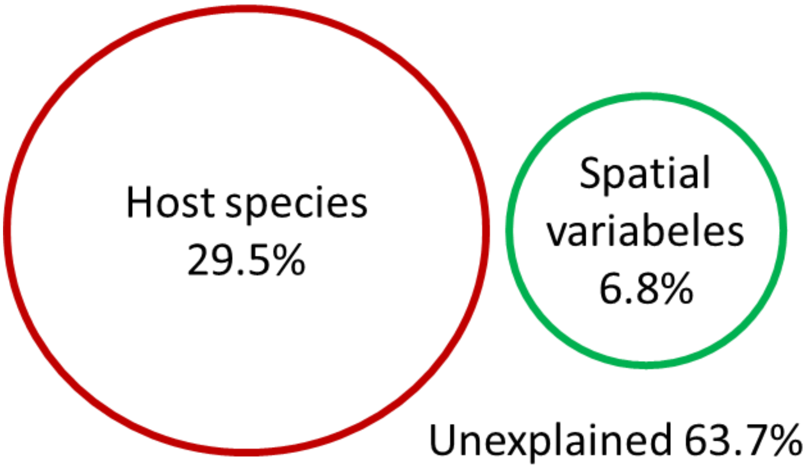


Fig. 3 Venn diagram showing the effects of host species and spatial distance on the ectomycorrhizal (ECM) fungal community composition as derived from the variation partitioning analysis using the Raup-Crick index. Numbers indicate the proportions of explained variation. No shared fraction between the host species and spatial variables was detected.

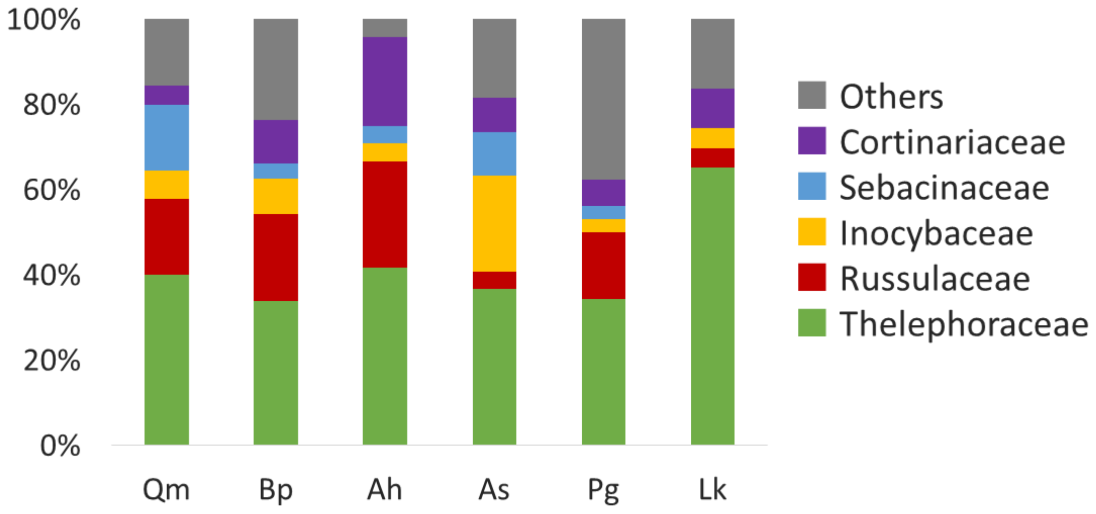


Fig. S4 Family level proportions of ECM fungal OTU for each host tree species.
